# Evaluating the clinical validity of gene-disease associations: an evidence-based framework developed by the Clinical Genome Resource

**DOI:** 10.1101/111039

**Authors:** Natasha T. Strande, Erin Rooney Riggs, Adam H. Buchanan, Ozge Ceyhan-Birsoy, Marina DiStefano, Selina S. Dwight, Jenny Goldstein, Rajarshi Ghosh, Bryce A. Seifert, Tam P. Sneddon, Matt W. Wright, Laura V. Milko, J. Michael Cherry, Monica A. Giovanni, Michael F. Murray, Julianne M. O’Daniel, Erin M. Ramos, Avni B. Santani, Alan F. Scott, Sharon E. Plon, Heidi L. Rehm, Christa L. Martin, Jonathan S. Berg

## Abstract

With advances in genomic sequencing technology, the number of reported gene-disease relationships has rapidly expanded. However, the evidence supporting these claims varies widely, confounding accurate evaluation of genomic variation in a clinical setting. Despite the critical need to differentiate clinically valid relationships from less well-substantiated relationships, standard guidelines for such evaluation do not currently exist. The NIH-funded Clinical Genome Resource (ClinGen) has developed a framework to define and evaluate the clinical validity of gene-disease pairs across a variety of Mendelian disorders. In this manuscript we describe a proposed framework to evaluate relevant genetic and experimental evidence supporting or contradicting a gene-disease relationship, and the subsequent validation of this framework using a set of representative gene-disease pairs. The framework provides a semi-quantitative measurement for the strength of evidence of a gene-disease relationship which correlates to a qualitative classification: “Definitive”, “Strong”, “Moderate”, “Limited”, “No Reported Evidence” or “Conflicting Evidence.” Within the ClinGen structure, classifications derived using this framework are reviewed and confirmed or adjusted based on clinical expertise of appropriate disease experts. Detailed guidance for utilizing this framework and access to the curation interface is available on our website. This evidence-based, systematic method to assess the strength of gene-disease relationships will facilitate more knowledgeable utilization of genomic variants in clinical and research settings.

## INTRODUCTION

The human genome comprises approximately 20,000 protein-coding genes^1^, of which about 3,000 have been reported in association with at least one Mendelian disease^2^. Roughly half^2^ of these gene-disease relationships have been identified over the last decade, as technological advances have made it possible to use sequence information from small families or even single individuals to discover new candidate gene-disease relationships^3; 4^ However, there is substantial variability in the level of evidence supporting these claims, and a systematic method for curating and assessing evidence is needed.

Despite this variability, clinical laboratories may include genes with preliminary evidence of a gene-disease relationship on disease-targeted panels, or in results returned from exome/genome sequencing. Some of the gene-disease relationships are either unable to be confirmed for many years or are ultimately proven wrong^5^. Evaluating the clinical impact of variants identified in genes with an unclear role in disease is exceedingly difficult, and could lead to incorrect diagnoses, preventing further evaluations and/or resulting in errant management of the affected individual and their families. This scenario highlights the need for a standardized method to evaluate the evidence implicating a gene in disease and thereby determine the clinical validity^3^ of a gene-disease relationship.

The NIH-funded Clinical Genome Resource (ClinGen)^6^ is creating an open-access resource to better define clinically relevant genes and variants based on standardized, transparent evidence assessment for use in precision medicine and research. Our group has developed a method that 1) qualitatively defines gene-disease clinical validity using a classification scheme based on the strength of evidence supporting the relationship, and 2) provides a standardized semi-quantitative approach to evaluate available evidence and arrive at such a classification. Currently, this framework is optimized for genes associated with monogenic disorders following autosomal dominant, autosomal recessive, or X-linked inheritance. Future iterations will expand the framework to consider other modes of inheritance, such as mitochondrial, and diseases with more complex genomic etiologies, including oligogenic or multifactorial conditions. Our approach is neither intended to define multifactorial disease risk, nor to be a substitute for well-established statistical thresholds used for genome-wide association studies^7; 8^.

This novel framework classifies gene-disease relationships by the quantity and quality of the evidence supporting such a relationship. It builds on efforts to catalog gene-disease associations, such as the Online Mendelian Inheritance in Man (OMIM)^1^ and OrphaNet, by systematically organizing the supporting and refuting evidence, and categorizing the strength of evidence supporting these relationships. The resulting clinical validity classifications are valuable to both clinicians and clinical laboratories. First, they provide insight into the strength of clinical associations for clinicians interpreting genetic test results for clinical care. Second, they serve to guide clinical genetic testing laboratories as they develop disease-specific clinical genetic testing panels or interpret genome-scale sequencing tests. By including only those genes with established clinical validity, the possibility of returning ambiguous, incorrect, or uninformative results is reduced, improving the quality of interpretation of genomic data.

## QUALITATIVE DESCRIPTION: CLINICAL VALIDITY CLASSIFICATIONS

The ClinGen Gene Curation Working Group (GCWG) is comprised of medical geneticists, clinical laboratory diagnosticians, genetic counselors, and biocurators with broad experience in both clinical and laboratory genetics. Over the course of three years, this group convened bi-monthly to develop the described framework for assessing gene-disease clinical validity through expert opinion and working group consensus (additional details provided on the ClinGen website). We first defined six classes to qualitatively describe the strength of evidence supporting a gene-disease association (Figure 1). The amount and type of evidence required for each clinical validity classification builds upon that of the previous classification level. Evidence used within this framework to assign a classification to a gene-disease pair is divided into two main types: genetic evidence and experimental evidence (described below). As evidence is likely to change over time, any given classification is only representative of the level of evidence at the time of curation.

**Figure 1:**
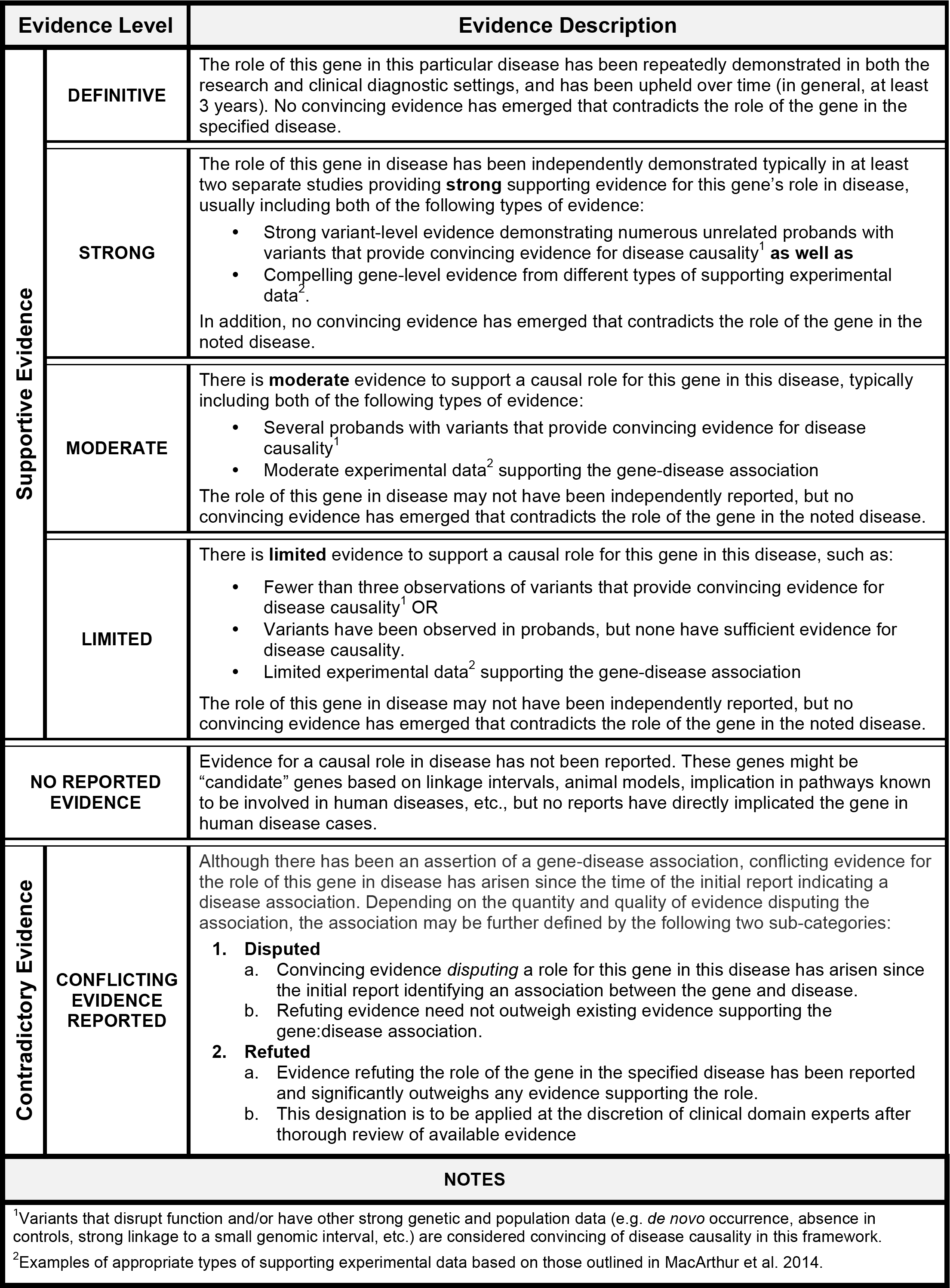
ClinGen clinical validity classifications and qualitative descriptions. The suggested minimum criteria needed to obtain a given classification are described for each clinical validity classification. The types of evidence comprising these criteria are described in the text. The default classification for genes without a convincing human disease-causing variant is “No Reported Evidence.” The level of evidence needed for each supportive gene-disease association category builds upon the previous category (i.e. “Limited” builds upon “Moderate”). Gene-disease associations classified as “Contradictory” likely have supporting evidence as well as opposing evidence, but are described separately from the classifications for supportive gene-disease associations.

The classification “No Reported Evidence” is used for genes that have not yet been asserted to have a causal relationship with a human monogenic disorder, but may have some experimental data (e.g., model system data) suggesting a potential role for that gene in disease. The “Limited” classification requires at least one variant, asserted to be disease-causing, to have plausible genetic evidence to support the association with human disease with or without gene-level experimental data. “Moderate” classification encompasses additional clinical evidence (e.g. multiple unrelated probands harboring variants with potential roles in disease) and supporting experimental evidence, all of which may be provided by multiple studies or a single robust study. Replication of the gene-disease association in subsequent independent publications and additional substantial genetic and experimental data are critical factors for the “Strong” classification. Finally, the hallmark of a “Definitive” gene-disease association is that, in addition to the accumulation of convincing genetic and experimental evidence, the relationship has been replicated, and ample time has passed since the initial publication (in general, greater than three years) for any conflicting evidence to emerge. It is important to highlight that these classifications do not reflect the effect size or relative risk attributable to variants in a particular gene, but instead the strength of the evidence. For example, a definitive gene-disease association does not imply that a pathogenic variant in that gene confers 100% penetrance of the phenotype. This metric is not intended to assess the penetrance or risk to develop a disease outcome.

A gene-disease relationship can be determined to have one of the above classifications provided no substantial relevant and valid contradictory evidence exists to call the gene-disease relationship into question. If such evidence emerges, then the relationship is described as “Conflicting Evidence Reported.” Types of contradictory evidence may come from population studies (such as ExAC^9^), attempts to experimentally validate the gene-disease association, or re-analysis of the original family or cohort that was previously studied. Although the role of a specific *variant* in a given disease may be called into question by new evidence, this may not be sufficient to invalidate the role of the *gene* in that disease. Thorough evaluation by experts in the particular disease area is recommended to determine whether the contradictory evidence outweighs the existing supportive evidence to classify a gene into either a “Disputed” or “Refuted” category (see Figure 1 for additional details).

## METHODS: SEMI-QUANTITATIVE ASSESSMENT OF EVIDENCE

Assigning a clinical validity classification to a gene-disease pair requires assessment of the evidence supporting the association. We developed a semi-quantitative approach to evaluate both genetic (Figure 2) and experimental evidence (Figure 3) in a standardized manner that promotes consistent collection and weighting of evidence (a detailed standard operating procedure is available on the ClinGen website). Defined sub-categories of genetic and experimental evidence are given a suggested default “score.” However, given that evidence of the same general type may vary in its strength (particularly when considering different diseases), the scoring system also allows these scores to be adjusted within a set range of points, with final approval by experts within the particular disease domain. Finally, the maximum number of points allowed for the various types of genetic and experimental evidence is capped to prevent a preponderance of weak evidence from inappropriately inflating the gene-disease classification. Similarly, certain evidence categories are provided higher maximum scores, allowing key pieces of stronger evidence to proportionately influence the classification of a gene-disease pair.

**Figure 2:**
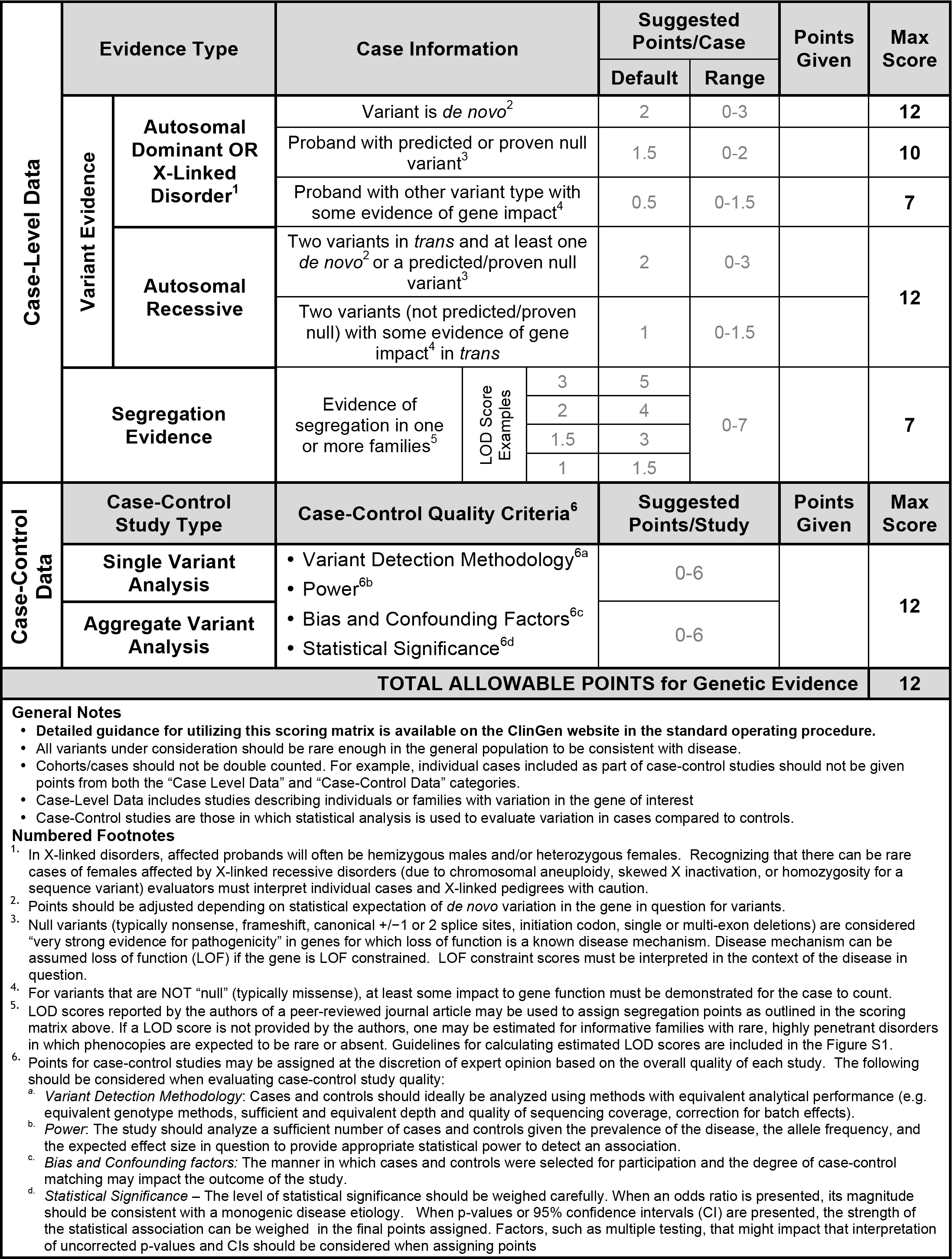
Classes of genetic evidence and their relative weights used in the ClinGen clinical validity framework. For additional points to consider when scoring genetic evidence, please see the standard operating procedure document available on our website. Genetic evidence is separated into two main categories: case-level data and case-control data. While a single publication may include both case-level and case-control data, individual cases should NOT be included in both categories. Each category is assigned a range of points with a maximum score that can be achieved. *<u>Case-Level Data</u>* is derived from studies describing individuals and/or families with qualifying variants in the gene of interest. Points should be assigned to each case based on the variant’s inheritance pattern, molecular consequence and evidence of pathogenicity in disease. In addition to variant evidence points, a gene-disease pair may also receive points for compelling segregation analysis (see Figure S1). *<u>Case-Control Data:</u>* Studies utilizing statistical analysis to evaluate variants in cases compared to controls. Case-control studies can be classified as either single variant analysis or aggregate variant analysis, however the number of points allowable for either category is the same. Points should be assigned according to the overall quality of each study based on these criteria: variant detection methodology, power, bias and confounding factors, and statistical power. Note that the maximum total scores allowed for different types of Case-Level data are not intended to add up to the total points allowed for Genetic Evidence as a whole. This permits different combinations of evidence types to achieve the maximum total score.

**Figure 3:**
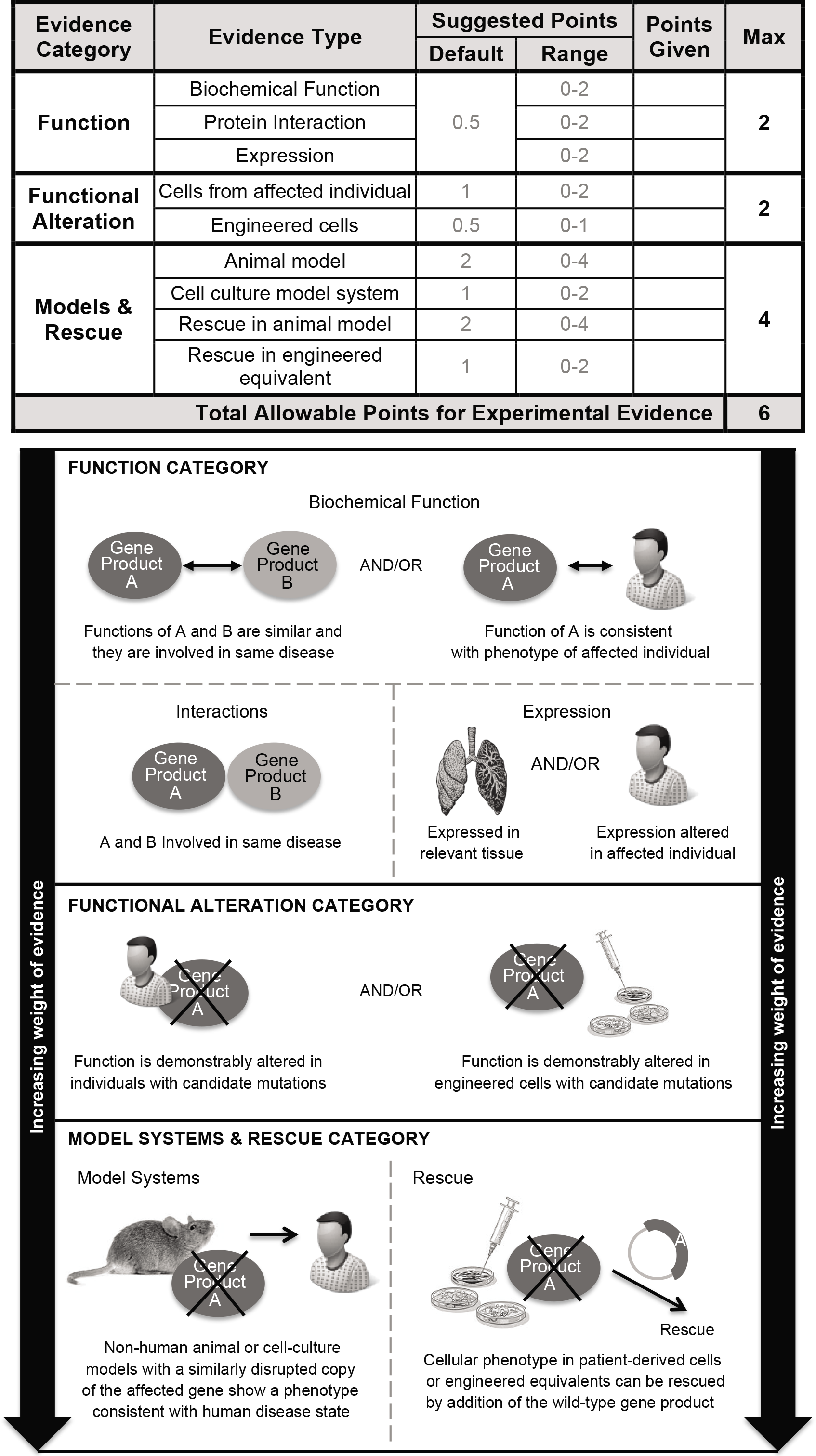
Types of gene-level experimental evidence and their relative weights used in the ClinGen clinical validity framework. Experimental evidence types used in the ClinGen gene curation framework are modified from MacArthur, et al. 2014. Evidence types are divided into three categories based on their relative contribution to the overall clinical validity of a gene-disease pair giving more weight to *in vivo* data. Each category is assigned a range of points with a maximum score that can be achieved, allowing more weight to be given to *in vivo* data (e.g. Models & Rescue) over *in vitro* experimental data. Evidence within the function category is given the least weight and is comprised of the following types of evidence: biochemical function, interactions, and expression. Functional alteration experiments in cells from affected individuals carrying candidate pathogenic variants are given more weight than the function category. Finally, model systems and phenotypic rescue experiments are given the most weight in our framework. Note that the maximum total scores allowed for different categories of Experimental Evidence are not intended to add up to the total allowable points. This permits different combinations of evidence types to achieve the maximum total score.

### Genetic Evidence

For the purposes of scoring, genetic evidence is divided into two categories: case-level data and case-control data (Figure 2). Studies describing individuals or families with genetic variants are scored as case-level data, while studies using statistical analyses to compare variants in cases and controls are scored as case-control data. When case-level and case-control data are present in a single publication, points can be assigned in each category, but the same piece of evidence should not be counted more than once. For example, an individual case that is also included within a case-control cohort should not be given points in both the “case-level data” and “case-control data” categories. In this scenario, points should be assigned to the most compelling and informative evidence.

Assessing case-level data requires consideration of the inheritance pattern and evaluation of the individual variants identified in each case. Within this framework, a case should only be counted towards supporting evidence if the reported variant has some indication of a potential role in disease (e.g., impact on gene function, recurrence in affected individuals, etc.), does not have evidence that would contradict pathogenicity (e.g., population allele frequency), and is of the type consistent with the assumed disease mechanism (e.g. truncating variant for loss of function). Unless otherwise noted, the term “qualifying variant” implies that these criteria are met. In addition, points are assigned separately for segregation data to reflect the statistical probability that the locus is implicated in the disease. Figure 2 and Figure S1 provide guidance on the number of points that should be considered for segregation evidence by LOD score; if a LOD score is not provided within the publication being evaluated, an estimated LOD score may be calculated in certain scenarios, as described in the standard operating procedure document provided on the ClinGen website.

Each study categorized as “case-control data” should be independently assessed to evaluate the quality of the study design (see Figure 2). Consultation with a clinical domain expert group (such as those affiliated with ClinGen, https://www.clinicalgenome.org/working-groups/clinical-domain/) is recommended. For the purposes of this framework, studies are classified based on whether they include single variant analysis or aggregate variant analysis. Single variant analyses are those in which individual variants are evaluated for statistical enrichment in cases compared to controls. More than one variant may be analyzed, but the variants have been independently assessed with appropriate statistical correction for multiple testing. Aggregate variant analyses are those in which the total number of variants is assessed for enrichment in cases compared with controls. This comparison is typically accomplished by sequencing the entire gene in both cases and controls and demonstrating an increased “burden” of variants of one or more types.

### Experimental Evidence

The experimental data scoring system is presented in Figure 3. The gene-level experimental data used in this framework to assess a gene-disease association are consistent with those proposed by MacArthur and colleagues to implicate a gene in disease^10^. The following experimental evidence types are used: biochemical function, experimental protein interactions, expression, functional alteration, phenotypic rescue and model systems (Figure 3 bottom panel). These categories capture the most relevant types of experimental information necessary to determine whether the function of the gene product is at least consistent with the disease with which it is associated, if not causally implicated.

### Contradictory Evidence

While curators are encouraged to seek out and document (via qualitative description) conflicting evidence, no specific points are assigned to this category. The types of valid contradictory evidence and their relative weights will be unique to each gene-disease pair, and it would be misleading to attempt to uniformly quantify this type of negative evidence against the reported positive evidence. If there is substantial conflicting evidence, manual review and expert input is required to evaluate the strength of the contradictory evidence, determine whether it outweighs any available supporting evidence, and, if so, decide whether the gene-disease association should be classified as “Disputed” or “Refuted”.

### Summary & Final Matrix

The scores assigned to both genetic and experimental evidence are tallied to generate a total score (ranging from 1-18) that corresponds to a preliminary clinical validity classification (Figure 4). The system provides a transparent method for summarizing and assessing all curated evidence for a gene-disease pair, encouraging consistency between curators. While the summary matrix facilitates a preliminary assessment of the gene-disease relationship, the initial curator or expert reviewer may adjust the classification, supplying a specific rationale for the change. Final classifications are determined in collaboration with disease experts, who review the preliminary classification and supporting evidence and work to come to a consensus with the preliminary curators. In the event that the disease experts and preliminary curators disagree on a final classification, a senior member of the ClinGen Gene Curation Working Group may be brought in to facilitate a final classification, erring towards the more conservative classification if consensus cannot be achieved. It should be noted that experimental data alone cannot justify a clinical validity classification beyond “No Reported Evidence,” and at least one human genetic variant with a plausible causal association must be present to attain “Limited” classification. The difference between “Limited,” “Moderate,” and “Strong” gene-disease classifications is justified by the quality and quantity of evidence; it is expected that valid gene-disease associations will gradually accumulate enough supporting evidence and be replicated over time to attain a “definitive” classification. This framework relies predominantly on evidence obtained from published primary literature, identified through resources such as PubMed and OMIM^1^, and independently assessed by curators; however, if necessary, unpublished information available from publicly accessible resources, such as variant databases^11; 12^, may be used as long as some supporting evidence is provided.

**Figure 4.**
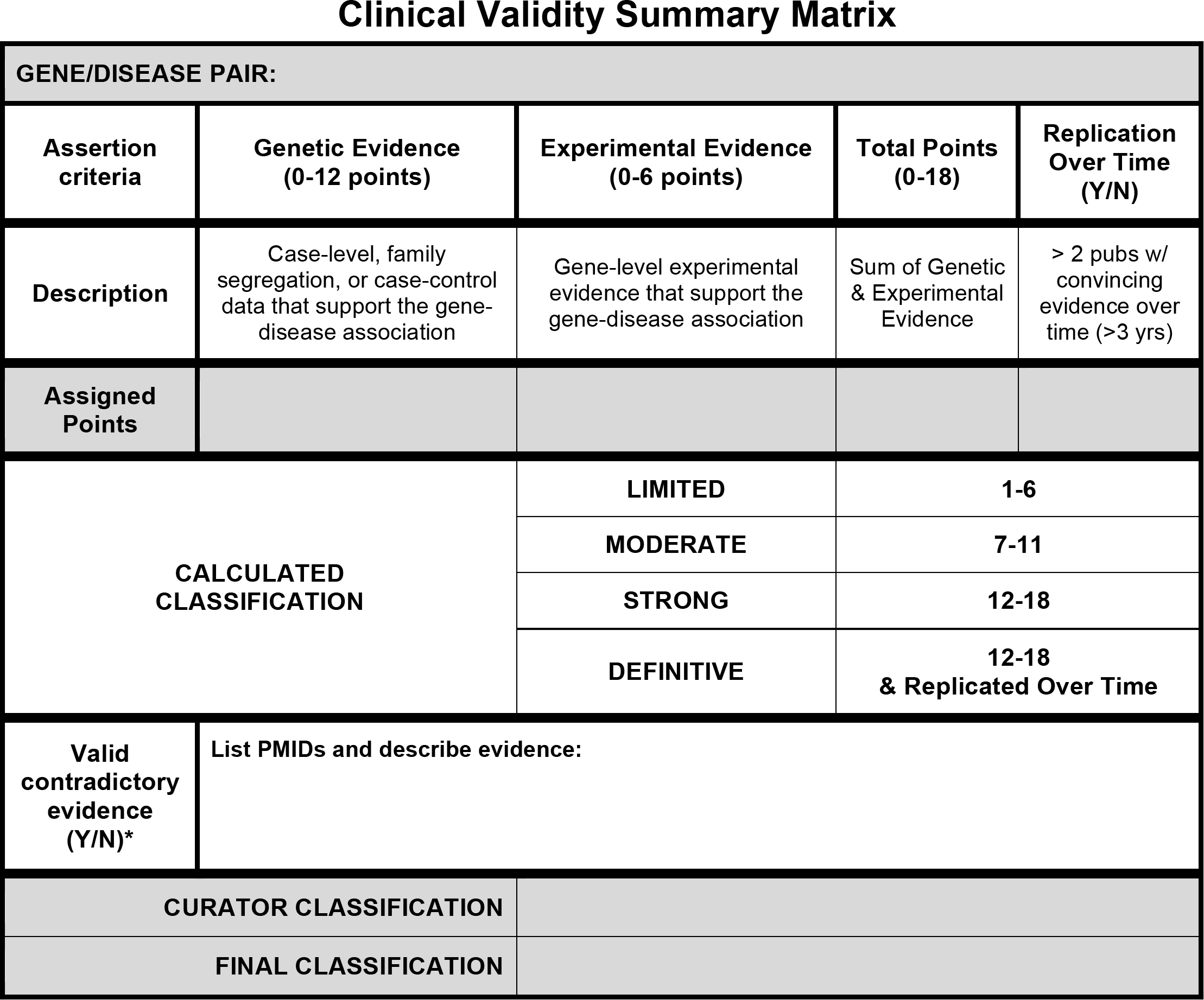
Final summary matrix used to provisionally classify gene-disease associations. A summary matrix was designed to generate a “provisional” clinical validity assessment using a point system consistent with the qualitative descriptions of each classification. *<u>Genetic Evidence:</u>* total number of points (not exceeding 12) obtained using the scoring metric in Fig. 2. If no human variants associated with disease have been reported in the literature, then the default classification is “No Reported Evidence.” *<u>Experimental Evidence</u>:* total number of points (not exceeding 6) derived from each of the experimental categories in Fig. 3. *<u>Replication Over Time</u>* - Yes, if more than three years has passed since the publication of the first paper reporting the gene-disease relationship AND more than two publications with human mutations exist. *<u>Contradictory Evidence</u>* - No points are assigned to this category. Instead, the curator should provide a summary of contradictory information. *<u>Scoring</u>* - The sum of the quantified evidence from each category can be used to determine a “provisional” classification using the scale at the bottom of the figure. If a curator does not agree with this classification, he/she may provide a different suggested classification along with appropriate justification.

## RESULTS: VALIDATION OF METHOD

Using this framework we evaluated 33 gene-disease pairs representing a variety of disease domains and spanning the spectrum of clinical validity classifications (see Table 1, Figure 5, and Supplemental Appendix). To assess the reproducibility of our scoring metric, each gene-disease pair was evaluated by two independent curators; paired curators reached concordant clinical validity classifications in 29 of the 31 (93.5%) gene-disease pairs with available published evidence (Figure 5; associations classified as “No Reported Evidence” were excluded). Each gene-disease pair was subsequently reviewed by clinical domain experts; experts agreed with the preliminary classifications for 87.1% (27/31) of the gene-disease pairs with published evidence (Figure 5). The four discrepancies between the expert and curator classifications were each different by only a single category (e.g. limited versus moderate). Of note, the original classifications for *HNRNPK* (MIM 600712) and *SMARCA1* (MIM 300012) were at the border between limited and moderate (6.5 points); in each case, the preliminary curators’ lack of specific clinical expertise led to uncertainty regarding the scoring of evidence requiring such knowledge. Consulting with clinical experts in the disease resolved these issues resulting in both genes being upgraded to moderate. In the case of *WRAP53* (MIM 612661), the expert was aware of additional published experimental evidence that when included increased the classification from limited to moderate. Upon reviewing the curated evidence for *RAD51D* (MIM 602954) and breast cancer (MIM 614291), the domain expert upgraded the classification from disputed to limited (with the approval of the GCWG) due to the specificity of the experimental evidence and insufficient power of the current studies to rule out a role for *RAD51D* in breast cancer (Figure 5). Details and references for each curation are provided in Supplemental Appendix.

**Table 1:** Categorization of gene-disease pairs used to validate the gene-validity framework

| Disease category | HGNC Gene Symbol | Gene MIM ID | Disease curated | Inheritance Pattern | Orphanet ID/ Phenotype MIM ID | Expert Reviewed Classification<sup>a</sup> |
| --- | --- | --- | --- | --- | --- | --- |
| Bone Marrow Failure | *NHP2* | 606470 | Dyskeratosis congenita | Recessive | ORPHA1775/MIM 613987 | Limited |
|  | *RAD51C* | 602774 | Fanconi anemia | Recessive | ORPHA84/MIM 613390 | Moderate |
|  | *RPS10* | 603632 | Diamond-Blackfan anemia | Dominant | ORPHA124/MIM 613308 | Definitive |
|  | *RPS24* | 602412 | Diamond-Blackfan anemia | Dominant | ORPHA124/MIM 610629 | Definitive |
|  | *TSR2* | 300945 | Diamond-Blackfan anemia with mandibulofacial dysostosis | X-Linked | ORPHA124/MIM 300946 | Limited |
|  | *WRAP53* | 612661 | Dyskeratosis congenita | Recessive | ORPHA1775/MIM 613988 | Moderate |
| Cardiovascular Disorders | *AKAP9* | 604001 | Romano-Ward syndrome | Dominant | ORPHA101016/MIM 611820 | Limited |
|  | *SCN4B* | 608256 | Long QT syndrome | Dominant | ORPHA768/MIM 611819 | Limited |
|  | *SMAD3* | 603109 | Loeys-Dietz type 3 | Dominant | ORPHA284984/MIM 613795 | Definitive |
|  | *TMPO* | 188380 | Familial or idiopathic dilated cardiomyopathy | Dominant | ORPHA154/MIM 613740<sup>b</sup> | Contradictory (refuted) |
| Hereditary Cancer | *DICER1* | 606241 | Pleuropulmonary blastoma | Dominant | ORPHA64742/MIM 601200 | Definitive |
|  | *PALB2* | 610355 | Hereditary breast cancer | Dominant | ORPHA227535/MIM 114480 | Definitive |
|  | *PMS2* | 600259 | Hereditary pancreatic cancer | N/A | N/A | No Reported Evidence |
|  | *RAD51D* | 602954 | Hereditary breast cancer | Dominant | ORPHA227535/MIM 614291 | Limited |
| Immune Disorders | *C1QB* | 120570 | Immunodeficiency due to C1Q deficiency | Recessive | ORPHA169147/MIM 613652 | Definitive |
|  | *CD3E* | 186830 | Severe combined immunodeficiency | Recessive | ORPHA183660/MIM 615615 | Definitive |
| Skeletal Dysplasia | *ARSD* | 300002 | Chondrodysplasia punctata | N/A | N/A | No Reported Evidence |
|  | *COL2A1* | 120140 | Spondyloepiphyseal dysplasia (Stanescu type) | Dominant | ORPHA94068/MIM 616583 | Moderate |
|  | *FGFR3* | 134934 | Achondroplasia | Dominant | ORPHA15/MIM 100800 | Definitive |
|  | *LBR* | 600024 | Anadysplasia-like, spontaneously remitting spondylometaphyseal dysplasia | Recessive | ORPHA448267/None | Moderate |
| Neuromuscular Disorders | *BAG3* | 603883 | Myofibrillar myopathy | Dominant | ORPHA593/MIM 612954 | Definitive |
|  | *MYO9A* | 604875 | Arthrogryposis | Recessive | ORPHA109007/None | Limited |
|  | *PSD3* | 614440 | Antecubital pterygium syndrome | Dominant | ORPHA2987/None | Limited |
|  | *VPS8* | N/A | Arthrogryposis | Recessive | ORPHA109007/None | Limited |
| Miscellaneous | *AGTR2* | 300034 | X-linked non-syndromic intellectual disability | X-Linked | ORPHA777/None | Contradictory (Disputed) |
|  | *ATF6* | 605537 | Achromatopsia | Recessive | ORPHA49382/MIM 616517 | Strong |
|  | *CHD1L* | 613039 | Renal or urinary tract malformation | Dominant | ORPHA93545/None | Limited |
|  | *HNRNPK* | 600712 | Au-Kline syndrome | Dominant | ORPHA453504 /MIM 616580 | Moderate |
|  | *LAMB1* | 150240 | Lissencephaly 5 | Recessive | ORPHA352682/MIM 615191 | Moderate |
|  | *NGLY1* | 610661 | Congenital disorder of deglycosylation | Recessive | ORPHA404454/MIM 615273 | Definitive |
|  | *SMARCA1* | 300012 | Syndromic intellectual disability with Coffin-Syris-like features | Dominant | None/None | Moderate |
|  | *SKI* | 164780 | Shprintzen-Goldberg | Dominant | ORPHA311140/MIM 182212 | Definitive |
|  | *SOS2* | 601247 | Noonan syndrome | Dominant | ORPHA648/MIM 616559 | Moderate |

<sup>a</sup> All gene-disease classifications are accurate as of January 2017.
<sup>b</sup> Phenotype MIM was associated with *TMPO* at the time of curation, but has since been removed due to updated information.
Abbreviations: N/A, not applicable.

**Figure 5.**
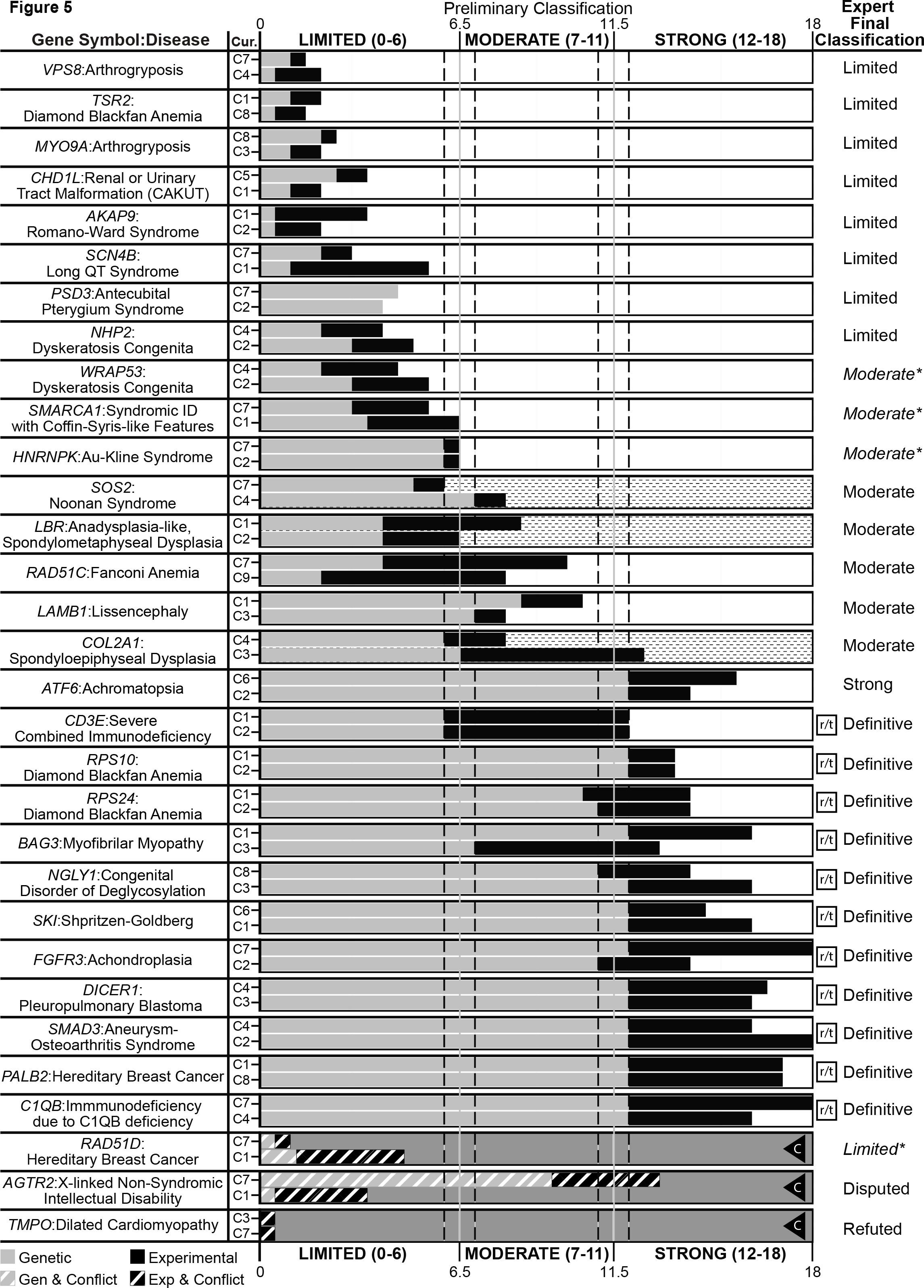
Comparison of provisional clinical validity classifications and associated matrix scores for selected gene-disease pairs evaluated by multiple curators. Of the 33 gene-disease pairs (y-axis) curated to validate the clinical validity curation framework, 31 were classified using the summary matrix (2 gene-disease pairs, *PMS2*:pancreatic cancer and /*ARSD*:chondrodysplasia punctata, were classified as “No evidence reported” and are not shown). Genetic evidence (grey bars) and experimental evidence (black bars) were evaluated by two independent curators (C1-C9) to arrive at a provisional classification (x-axis). Gene-disease relationships scoring between 12-18 points can be “Strong” or “Definitive,” depending on whether the association has been replicated over time (indicated by the squared “r/t”), in which case the preliminary classification is “Definitive”. Clinical validity classifications that were discordant between preliminary curators are represented with a dashed background. Gene-disease pairs in which conflicting evidence was reported are represented by diagonal lines through the evidence bars and a grey background. The letter “C” in a triangle indicates that the curators classified the gene-disease pair as “Conflicting Evidence Reported”. Each gene-disease pair was ultimately evaluated by an expert in the field for a final classification (far right column). Final expert classifications that differed from the preliminary classification are indicated by italics and asterisks.

## DISCUSSION

The evidence-based framework described here qualitatively defines clinical validity classifications for gene-disease associations in monogenic conditions and provides a systematic framework for evaluating key criteria required for these classifications. This method is intentionally flexible to accommodate curation of a wide spectrum of genes and conditions by curators with varying levels of expertise. The semi-quantitative scoring system combined with the qualitative classification scheme guides curators through the preliminary decision-making process, while the expert-level review provides disease-specific experience to weigh in on the final classification.

This effort to create a generalized framework may result in some specific challenges due to the heterogeneity of genetic conditions, in both phenotype and prevalence. For example, conditions that span a large phenotypic spectrum may pose a challenge when defining what constitutes a condition and what is most relevant for curation purposes. In general, ClinGen encourages its expert curation groups to focus on disease associations that have been asserted in the literature or in other authoritative sources (e.g. OMIM, Orphanet Disease Ontology). Expert reviewers may find it useful in certain scenarios, to curate both a syndromic disease association as well as an isolated/non-syndromic disease association limited to a particular sub-phenotype. For example, when a disease entity encompasses sub-phenotypes that are caused by different mutational mechanisms. This is a topic of continued discourse within the ClinGen working groups and will be incorporated into future manuscripts that will focus on the curation approach for individual ClinGen disease-focused expert groups.

Ultra-rare disorders may have a relatively small number of probands described in the medical literature, thus limiting their potential to achieve a high genetic evidence score within this matrix. This obstacle is mostly circumvented by allowing compelling pieces of genetic evidence to score the maximum number of points (for example, see *CD3E* (MIM 186830) and severe combined immunodeficiency (MIM 615615), detailed in the Supplemental Appendix). When substantial experimental evidence is also available, these conditions can attain a “Strong” or “Definitive” classification. On the opposite end of the spectrum are conditions that occur commonly in the general population, such as cancer, where the predominant etiology is multifactorial rather than monogenic. In the less common Mendelian cancer predisposition syndromes, incomplete penetrance is a typical feature that can lead to confounding factors in family genetic studies such as apparently non-penetrant family members who carry a disease-associated variant and phenocopies among family members without a disease-associated variant. For such conditions, case-control data may provide more compelling evidence to support the gene-disease association (see the curation of *PALB2* (MIM 610355) and hereditary breast cancer (MIM 114480) in the Supplemental Appendix as an example).

One limitation of any such system is the challenge of balancing thorough literature curation and practical time commitment. This system can accommodate an exhaustive literature review, but in most cases will only require curating the amount of information sufficient to reach the maximum number of points in the matrix. In some scenarios this method may fail to include pertinent information, which could impact the classification (e.g., omission of contradictory evidence). Another potential limitation is the subjective nature of certain evidence types (e.g., experimental), which may lead to variability between different groups assessing evidence. However, due to the transparency of the evidence base, the incorporation of expert review, and the ability to reassess classifications over time, such drawbacks are likely to be self-limiting.

ClinGen’s ultimate goal is to enhance the incorporation of genomic information into clinical care, an important component of the Precision Medicine Initiative^13^. The implementation of this framework will be supported by an open-access ClinGen curation interface (under development) that will guide curators through the curation process and will serve as a platform for extension to the community. In essence, this framework aims to provide a systematic, transparent method to evaluate a gene-disease relationship in an efficient and consistent manner suitable for a diverse set of users. A detailed standard operating procedure for this framework is available on the ClinGen website. All curated evidence, including clinical validity assessments, will also be made readily accessible to clinical laboratories, clinicians, researchers, and the community via our website. Additionally, for community members that wish to contribute papers of interest and/or request curation of a gene-disease pair, a “reporter” form is available on the ClinGen website.

Carefully evaluated gene-disease clinical validity classifications, as provided by this framework, will be useful to clinical laboratories as they evaluate genes for inclusion on disease-targeted panels, or as they decide how to categorize, prioritize, and return results from exome/genome sequencing. Clinicians may choose to use these types of gene-disease classifications as they interpret laboratory results for the individuals they care for; for instance, they may choose not to adjust medical management based on variants in genes of limited clinical validity. Researchers could also utilize this framework to evaluate the clinical validity of their own newly discovered associations and identify promising target genes for future work in order to augment the currently available evidence and attain a “Strong” or “Definitive” classification. In addition, professional societies and regulatory bodies may utilize these clinical validity assessments when making recommendations or guidelines for clinical genetic testing. Ultimately, our systematic, evidence-based method for evaluating gene-disease associations will provide a strong foundation for genomic medicine.

## DESCRIPTION OF SUPPLEMENTAL DATA

The Supplemental file includes one figure, an appendix with curated evidence for each example presented in Figure 5, and a list of references. A more comprehensive supplemental file is available on the BioRxIV preprint server (see doi: https://doi.org/10.1101/111039)

## ACKNOWLEDGEMENTS

This work was supported by grants from the National Human Genome Research Institute (NHGRI), through the following three grants: U41 HG006834-01A1, U01 HG007437-01, U01 HG007436-01, as well as from the National Cancer Institute (NCI) through the following contract: HHSN261200800001E. ClinGen also receives funding through the Eunice Kennedy Shriver National Institute of Child Health and Human Development (NICHD) and ClinVar is supported by the Intramural Research Program of the NIH, National Library of Medicine. We would like to thank the following groups and individuals for contributing their disease expertise to review the examples included in this manuscript: Alan Beggs Ph.D.; Alison Bertuch, M.D., Ph.D.; Rebecca H. Buckley, M.D.; Eugene Chung, M.D.; Bill Craigen M.D., Ph.D.; Jennifer M. Puck, M.D.; Sharon A. Savage, M.D.; Fergus J. Couch Ph.D. and the ClinGen Hereditary Breast and Ovarian Cancer gene curation working group; Birgit H. Funke Ph.D. and the ClinGen Cardiomyopathy gene curation working group; and the ClinGen RASopathy curation working group. Input on the framework was also provided by the ClinGen Hereditary Breast and Ovarian Cancer gene curation working group and Ray Hershberger, Mike Gollob and the ClinGen Channelopathy gene curation working group. We would also like to thank Scott Goehringer for his invaluable help in preparing the curated examples for the ClinGen website and appendix.

## WEB RESOURCES

Clinical Genome Resource (ClinGen): www.clinicalgenome.org

ClinGen Gene Curation Working Group site: https://www.clinicalgenome.org/working-groups/gene-curation/

Standard operating procedure for the framework described in this manuscript: http://bit.ly/ClinGenGCSOP

Validation Method for the framework described in this manuscript: http://bit.ly/clingenGCValMethods

ClinGen “reporter” form: https://search.clinicalgenome.org/kb/agents/sign_up

OMIM: https://omim.org/

## REFERENCES

1. Online Mendelian Inheritance in Man, OMIM®. In. (Baltimore, MD, McKusick-Nathans Institute of Genetic Medicine, Johns Hopkins University

2. Chong, J.X., Buckingham, K.J., Jhangiani, S.N., Boehm, C., Sobreira, N., Smith, J.D., Harrell, T.M., McMillin, M.J., Wiszniewski, W., Gambin, T., et al. (2015). The Genetic Basis of Mendelian Phenotypes: Discoveries, Challenges, and Opportunities. American journal of human genetics 97, 199–215.

3. Haddow, J., Palomacki, G. (2003). ACCE: A Model Process for Evaluating Data on Emerging Genetic Tests. In Human Genome Epidemiology: A Scientific Foundation for Using Genetic Information to Improve Health and Prevent Disease, M. Khoury, Little, J., Burke, W., ed. (Oxford University Press), pp 217–233.

4. Wilfert, A.B., Chao, K.R., Kaushal, M., Jain, S., Zollner, S., Adams, D.R., and Conrad, D.F. (2016). Genome-wide significance testing of variation from single case exomes. Nat Genet 48, 1455–1461.

5. Eisenberger, T., Di Donato, N., Baig, S.M., Neuhaus, C., Beyer, A., Decker, E., Murbe, D., Decker, C., Bergmann, C., and Bolz, H.J. (2014). Targeted and genomewide NGS data disqualify mutations in MYO1A, the “DFNA48 gene”, as a cause of deafness. Human mutation 35, 565–570.

6. Rehm, H.L., Berg, J.S., Brooks, L.D., Bustamante, C.D., Evans, J.P., Landrum, M.J., Ledbetter, D.H., Maglott, D.R., Martin, C.L., Nussbaum, R.L., et al. (2015). ClinGen‐‐the Clinical Genome Resource. The New England journal of medicine 372, 2235–2242.

7. Lander, E., and Kruglyak, L. (1995). Genetic dissection of complex traits: guidelines for interpreting and reporting linkage results. Nat Genet 11, 241–247.

8. Sham, P.C., and Purcell, S.M. (2014). Statistical power and significance testing in large-scale genetic studies. Nature reviews Genetics 15, 335–346.

9. Lek, M., Karczewski, K.J., Minikel, E.V., Samocha, K.E., Banks, E., Fennell, T., O’Donnell-Luria, A.H., Ware, J.S., Hill, A.J., Cummings, B.B. et al., (2016). Analysis of protein-coding genetic variation in 60,706 humans. Nature 536, 285–291.

10. MacArthur, D.G., Manolio, T.A., Dimmock, D.P., Rehm, H.L., Shendure, J., Abecasis, G.R., Adams, D.R., Altman, R.B., Antonarakis, S.E., Ashley, E.A., et al. (2014). Guidelines for investigating causality of sequence variants in human disease. Nature 508, 469–476.

11. Landrum, M.J., Lee, J.M., Riley, G.R., Jang, W., Rubinstein, W.S., Church, D.M., and Maglott, D.R. (2014). ClinVar: public archive of relationships among sequence variation and human phenotype. Nucleic acids research 42, D980–985.

12. Fokkema, I.F., Taschner, P.E., Schaafsma, G.C., Celli, J., Laros, J.F., and den Dunnen, J.T. (2011). LOVD v.2.0: the next generation in gene variant databases. Human mutation 32, 557–563.

13. Collins, F.S., and Varmus, H. (2015). A new initiative on precision medicine. The New England journal of medicine 372, 793–795.

